# Stoichiometric analysis of microbial communities: interrelating community function, structure and biomass carrying capacity

**DOI:** 10.1101/2025.08.29.673032

**Authors:** Frank J Bruggeman, Timothy Paez-Watson, Bas Teusink, Robbert Kleerebezem

## Abstract

Microbial communities carry out important ecological functions. Their activities emerge from complex interactions between species, often potentiated by metabolic traits. We lack a quantitative understanding of how these traits shape community properties. Here, we present the theory for microbial communities, leveraging concepts from quantitative microbial physiology. We derive formal conditions for the steady states of microbial communities. We express the relative abundances of species (living and dead), the net metabolic conversion of a community, and the biomass carrying capacity in terms of the metabolic stoichiometry of the species and their growth and death rates. We show how niche creation can emerge from stoichiometric imbalances in cross-feeding communities. Finally, we discuss how relative species abundances depend on the ATP stoichiometries of intracellular metabolism.

## Introduction

Microbial communities play important roles in the planet-wide cycling of chemical elements, organismal health, and biotechnology (foods, natural fermentations, wastewater treatment) [1]. There is a need for a better understanding of these systems to forecast their responses to environmental changes (e.g. resilience and fragility) and to control their functions in applications. This is in particular relevant now, as many communities, and therewith their activities, are under threat of climate change and pollution (incl. antibiotics), and our economies need to make a shift to more bio-based production systems [2].

Metagenomics methods revolutionised our insight into the composition and genetic potential of the species making up communities [3, 4]. It remains challenging however to predict from this genomic knowledge emergent ecological mechanisms and activities of microbes in communities, e.g. how abundant they are, how fast they grow and die, what their nutrients and products are, and with whom they preferentially interact and why they do so [5, 6]. This prevents us from rationally steering the behaviour of communities.

Systems biology, biotechnology and quantitative microbial physiology have made great progress in our predictive understanding of the physiology of single microbes in controlled environments. This was primarily achieved by integrating theoretical concepts, simulations and quantitative data acquisition about, in particular, metabolic fluxes, yields, and (metabolic) protein concentrations [7, 8, 9]. For instance, we have currently an understanding of why and when microbes, at least for the model microbes *Escherichia coli, Saccharomyces cerevisiae* and *Synechocystis sp*., make switches in metabolic strategies across conditions (and growth rates) in general physiological terms such as optimal biosynthetic resource allocation, growth-rate maximisation, protein-expression constraints, and ATP yield on energy substrates [10, 11, 12]. We expect that these methods could also be adapted to microbiomes and thereby turned into powerful predictors of the behaviors of microbes in communities, given the dynamic environment that they partially generate themselves via ecological feedback [13]. Such an endeavor could effectively turn microbial ecology from a descriptive field into a more predictive discipline, focusing on control and steering of systems [2].

In this paper, we focus on the translation of yield prediction methods based stoichiometric and thermodynamic considerations, originally developed for studies of the physiology of single microbes [7, 8, 9], into a microbial ecology setting. Our methods allow for the prediction and understanding of the relative abundances of species and the net metabolic conversion and biomass carrying-capacity of the community, starting from the stoichiometric description of the metabolic activity and death rates of each of the species in the community. We limit this paper to the prediction of these systemic properties of the community at steady states. Steady states provide a natural starting point for understanding also more complex scenarios involving dynamics, since (stable) steady states serve as attractors in dynamic ecosystems [14]. In a follow-up study, we will focus on the dynamic description of microbial communities and these systemic quantities.

A central concept in this work is the macrochemical reaction (stating chemical-element conversion), a concept with a long history in quantitative microbial physiology and microbial ecology [7, 9]. One can think of the macrochemical reaction as the (chemical-element balanced) reaction converting substrates for growth and maintenance into biomass and byproducts, with a microbial species as the catalyst. A macrochemical reaction can be considered as a metabolic strategy of a microbe, e.g. it can grow via glucose respiration yielding carbon dioxide and water or via glucose fermentation giving rise to fermentation products. Microbes may shift between metabolic strategies when conditions change, affecting their product formation, cross-feeding interactions, and downstream metabolisms of other microbes [12]. Metabolic interactions therefore set the relative abundances of microbes in addition to their death processes. The entire set of macrochemical reactions that a microbe can manifest is a measure of its metabolic plasticity; nowadays, it can be computed from the microbe’s genome through metabolic network reconstruction methods [15, 16]. Macrochemical reactions can also be approximated from bioenergetic considerations. We refer to Kleerebezem et al. [17] for an overview of their derivations and applications in industrial applications.

Here we use macrochemical reactions of microorganisms to predict systemic stoichiometric properties of the community they compose, and subsequently derive the macrochemical reaction of the entire community (the community conversion) and determine the amount of biomass per unit energy source (the biomass carrying capacity).

## Results

### Macrochemical-equation based stoichiometric models of microbial communities

Macrochemical reactions result from the precise protein expression strategy of the species under the prevailing conditions and the assumption of a steady-state metabolism (balanced growth). Since during steady-state metabolism all intracellular metabolite concentrations remain constant, the chemical element composition of the growth substrates must balance with the composition of the growth products, including biomass. This elemental-conservation is captured in the macrochemical reaction, similarly as in any chemical reactions. The macrochemical equation can be calculated for any steady-state metabolic network from its stoichiometric matrix and steady-state flux distribution (see Appendix). Alternatively, a macrochemical equation can be deduced from experiments [7], estimated from chemical-element conservation and thermodynamic relations [8], and from genome-scale stoichiometric model of metabolism (Appendix; [18, 15, 19,16]).

An example of a macrochemical reaction for the anaerobic growth of *S. cerevisiae* on glucose with ethanol as the byproduct [7] is given by,

**Table 1.**
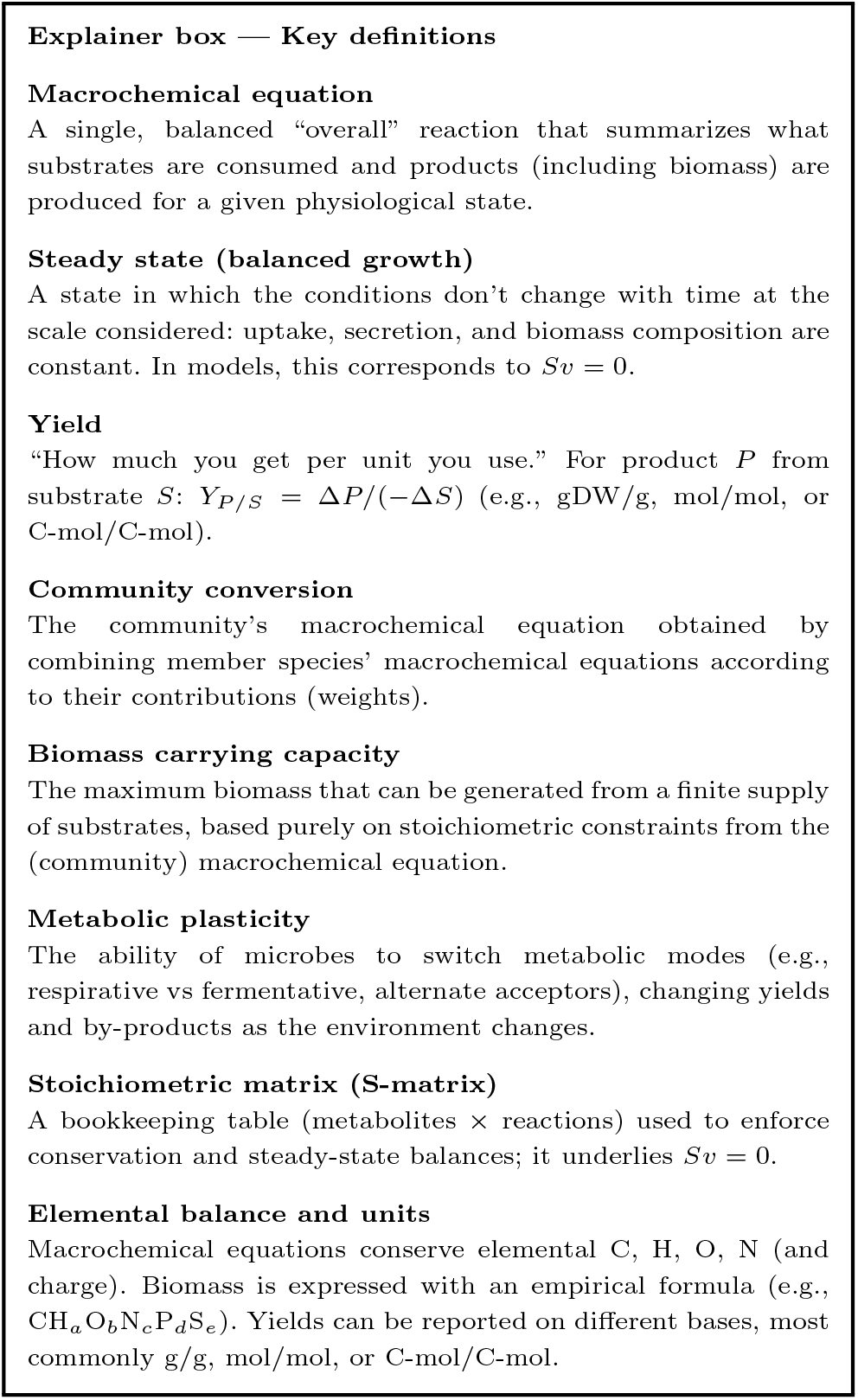
Explainer box with concise definitions for quick reference.

1. 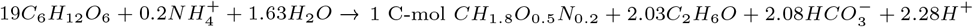

The coefficient in front of glucose (*C*_6_*H*_12_*O*_6_) specifies that 1.19 moles of glucose (denoted by G) is needed to make 1 C-mole of biomass (*CH*_1.8_*O*_0.5_*N*_0.2_; denoted by X) and 2.03 moles of ethanol (*C*_2_*H*_6_*O*; E) is made per C-mole of biomass. Thus, the yield (denoted by Y) of biomass on glucose equals *Y*_*X/G*_ = 1.19^*−*1^ C-mol/mol and the yield of ethanol on biomass equals *Y*_*E/X*_ = 2.03 mol/C-mol.

When the macrochemical reaction is written for 1 unit of biomass, the rate of the macrochemical reaction is equal to the specific growth rate (or per capita) (*µ* in C-mol/(C-mol hr)) of the associated species [9]. (We note that in some literature the specific growth rate is denoted by *q*_*X*_ or *λ*.) The amount of biomass can also be quantified in terms of gram dry weight instead of C-mole. The specific glucose uptake rate *q*_*G*_ (in mol glucose/(C-mol hr) and the specific ethanol production rate *q*_*E*_ (in mol ethanol/(C-mol hr)) are proportional to the growth rate, i.e. 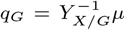 and *q*_*E*_ = *Y*_*E/X*_ *µ*. The proportionality constants are so called ‘yields’ denoted by *Y*_*P/S*_ for the yield of product *P* on substrate *S*. Similar proportionality relations hold between the other compounds’ specific rates and growth rate. The net flux of glucose uptake and ethanol production by yeast, when it occurs at a biomass abundance *X* (in C-moles or grams), equals *J*_*G*_ = *q*_*G*_*X* and *J*_*E*_ = *q*_*E*_ *X* respectively.

### A microbial community at steady state

Steady-state microbial communities are characterized by the constancy (time invariance) of: i. all concentrations of the extracellular metabolites, including those consumed and produced by the consortium members; ii. consequently, the growth rates and metabolic rates of the species, which depend on those concentrations; iii. the death rates of all the species, and iv. all intracellular concentrations (e.g., of proteins and metabolic intermediates; to achieve a state of “balanced growth”). Balanced growth and steady-state communities can be experimentally achieved under lab conditions (e.g., continuous cultures [20] and ‘approximated’ during the exponential phase in a batch cultivation [21]) and are approximated in applications, e.g., in wastewater treatment [22].

We define the relative abundance *ϕ*_*i*_ of a species *i* as its absolute abundance *X*_*i*_ (in C-mole or grams) divided by the total abundance of the community, 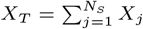, consisting of *N*_*S*_ species, i.e. *ϕ*_*i*_ = *X*_*i*_*/X*_*T*_.

When the community is in steady state, all the relative abundances of the species remain constant. This implies that their rate of change equal zero. In the appendix we derive that this rate of change equals,

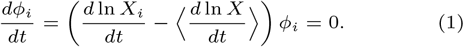

In this equation, 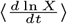 represents the average rate of change of the log abundance of all the species defined as 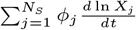 which equals *d* ln *X*_*T*_ */dt*, which we shown in the Appendix. Thus, at steady state, all species obey the following relations,

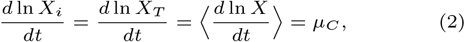

i.e., they all have the same net per-capita growth rate, which we shall refer to as the community growth rate *µ*_*C*_. This equation always applies, at any steady state, regardless of the occurrence of any other processes than growth that can alter the abundance of the species, such as their inflow (dispersion), outflow or death. Note that in many cases in nature, the net growth rate of a community may turn out be zero and the nutrient fluxes sustain both the maintenance requirements of the species and counter their death rates.

### The consideration of dead biomass

The consideration of the death rates of microorganisms forces the consideration of growing biomass (active) and dead biomass (inactive). Thus, the total biomass of a community or any of its species equals the sum of the associated living and dead biomass. We do not consider the recycling (growth on) dead biomass, since we currently lack any understanding of the stoichiometry of that process. This we see as an important open problem in the field [23].

We denote the abundance of the metabolically-active living biomass by ‘L’ and the dead biomass by ‘D’. The total abundance of species *i* therefore equals *X*_*i*_ = *L*_*i*_ + *D*_*i*_ and the total community biomass equals 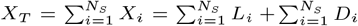.

We model the rates of change of the living, dead and total biomass of a species as,

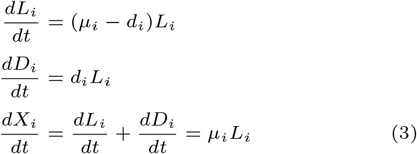

with *µ*_*i*_ and *d*_*i*_ respectively as the per-capita growth and death rate. Note that an extension of the theory, for instance, by incorporating other processes such as dispersion or predation would happen at this stage.

We introduce additional abundance fractions that can be distinguished: the relative abundance of the living and dead biomass of a species with respect to the total community biomass (i.e. *L*_*i*_*/X*_*T*_ and *D*_*i*_*/X*_*T*_ ) and with respect to the abundance of the species (i.e. *L*_*i*_*/X*_*i*_ and *D*_*i*_*/X*_*i*_). In the appendix, we show that the latter species-normalised abundances relate the growth rate and the death rates of the species in the following manner,

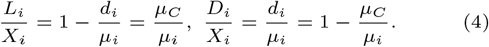

Thus, the death and living fraction of a species in a steady-state community can always be calculated from its total abundance using its growth rate and death rate. Note also that 0 *< L*_*i*_*/X*_*i*_ *<* 1 and 0 *< D*_*i*_*/X*_*i*_ *<* 1 such that 0 *< d*_*i*_*/µ*_*i*_ *<* 1.

From equations 2, 3 and 16, we conclude that *d* ln *X*_*i*_*/dt* = *µ*_*i*_ *− d*_*i*_ = *µ*_*C*_. This means that the death rate of a species can be determined from its (living and death) fractions and the community growth rate.

### Characteristics of a steady-state microbial community

Summarizing the results from the previous section, a steady-state microbial community is characterized by the following properties:

1. The growth rate of the entire community equals *d* ln *X*_*T*_ */dt* = *µ*_*C*_.
2. The concentrations of the exchanged nutrients and the physicochemical conditions are constant. Under these conditions, the growth and death rates of all species are also constant, and they satisfy *µ*_*C*_ = *µ*_*i*_ *− d*_*i*_ = *⟨µ − d⟩* (eq. 2).
3. Each species has a living and a dead fraction, given by *L*_*i*_*/X*_*i*_ = 1 *− µ*_*i*_*/d*_*i*_ = *µ*_*i*_*/µ*_*C*_ and *D*_*i*_*/X*_*i*_ = 1 *− L*_*i*_*/X*_*i*_.
4. The ratio of the living to dead abundance of a species equals *L*_*i*_*/D*_*i*_ = *µ*_*C*_ */d*_*i*_.

These relations are useful for determining the relative abundances of living and dead biomass from experimental data.

Without considering the macrochemical equations of growth for all species, it is not possible to determine the relative abundances *X*_*i*_*/X*_*T*_ (i.e., *ϕ*_*i*_). In the next sections, we address this for different microbial communities. We begin with a simple example, while the complete derivations are provided in the Appendix.

### The relative abundances of ethanol-exchanging yeast and acetobacter in a 2-species community

To illustrate the procedure for expressing the relative abundances of species in terms of the stoichiometric properties of their metabolisms (i.e., their macrochemical equations), we consider a 2-species community responsible for wine turning sour [24]. This community consists of a yeast producing ethanol from glucose and an acetobacter converting the ethanol into acetic acid (Figure 1A).

**Fig. 1.**
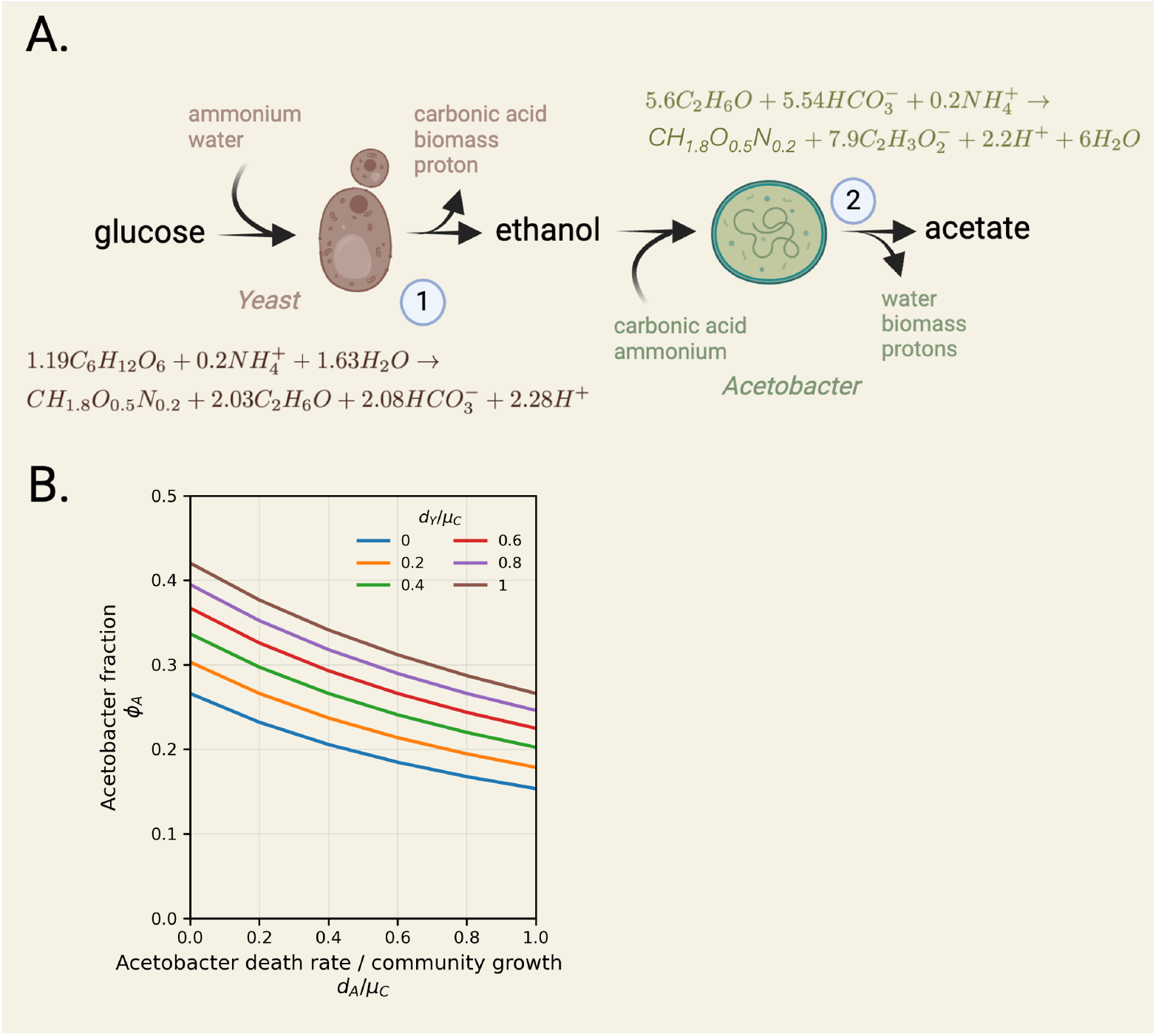
A 2-species community composed of a yeast and acetobacter converting glucose into acetate, via ethanol. A. The metabolic network formed by the cross-feeding of ethanol by a yeast and acetobacter strain is shown together with macrochemical equations. B. The relative abundances of acetobacter is shown as function of its relative death rate.

At steady state, the ethanol production rate by the yeast (*J*_*Y,E*_ ) equals the ethanol consumption rate by the acetobacter (*J*_*A,E*_ ), i.e., *J*_*Y,E*_ = *J*_*A,E*_. The net growth rates of both species equal the community growth rate:

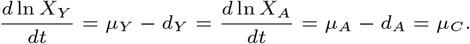

In a batch-cultivation scenario, the biomasses *X*_*Y*_, *X*_*A*_, and *X*_*T*_ would all increase exponentially with rate constant *µ*_*C*_. In a continuous culture (e.g., a chemostat), these rates would be set by the dilution rate, and biomass abundances would remain constant. The derivations that follow apply to all balanced-growth states of this community, regardless of the cultivation method.

The production rate *J*_*Y,E*_ of ethanol (*E*) by the yeast (*Y* ) is given by

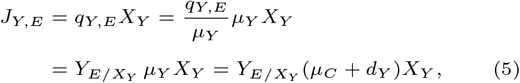

where 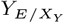 is the stoichiometric coefficient of ethanol in the yeast’s macrochemical growth equation (2.03 in Figure 1A).

Similarly, the consumption rate *J*_*A,E*_ of ethanol by the acetobacter (*A*) is

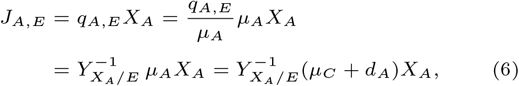

where 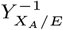 is the stoichiometric coefficient of ethanol (5.6 in Figure 1) in the macrochemical growth equation for acetobacter.

We could alternatively define *J*_*Y,E*_ = *q*_*Y,E*_ *L*_*Y*_ instead of *J*_*Y,E*_ = *q*_*Y,E*_ *X*_*Y*_, and similarly for acetobacter. This would require expressing the specific production and consumption rates in terms of living biomass only. However, because living biomass is often not directly measurable in experimental settings, we use total biomass here. This choice does not qualitatively change the following derivations.

At steady state, the ethanol production rate by the yeast balances the consumption rate by the acetobacter:

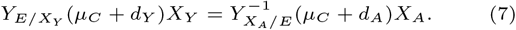

Dividing by the total abundance *X*_*T*_ = *X*_*A*_ + *X*_*Y*_ gives an equation in terms of the relative abundances *ϕ*_*Y*_ = *X*_*Y*_ */X*_*T*_ and *ϕ*_*A*_ = *X*_*A*_*/X*_*T*_. Since these sum to one (*ϕ*_*Y*_ + *ϕ*_*A*_ = 1), the expression can be solved for the relative abundance of either species. For acetobacter:

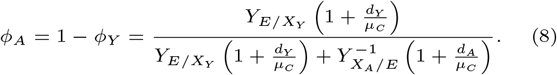

We can parameterise equation (8) using 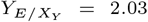 and 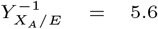, obtained from the macrochemical growth equations of the two microbes (Figure 1A). This parameterisation allows us to predict the relative abundance of acetobacter as a function of its death rate, normalised by the community growth rate (Figure 1B). The prediction aligns with intuition: the relative abundance of acetobacter decreases as its own death rate increases, and increases when the death rate of the yeast rises.

### Insights into the role of the death rate

Equation 8, or equivalently

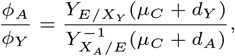

shows that, for a given community growth rate *µ*_*C*_, an increase in the acetobacter death rate *d*_*A*_ would require a higher acetobacter growth rate *µ*_*A*_ = *µ*_*C*_ + *d*_*A*_, yet its relative abundance would still decrease. Conversely, a higher yield of ethanol per unit yeast biomass (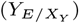) or a higher yield of acetobacter biomass per unit ethanol (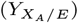) would increase the acetobacter fraction.

So far, we have considered communities with *µ*_*C*_ *>* 0, where *µ*_*i*_ *> d*_*i*_ for all species. In the special case *µ*_*C*_ = 0, the growth and death rates are equal for each species (*µ*_*i*_ = *d*_*i*_), leading to a stationary biomass despite ongoing metabolic activity.

### Growth conditions and nutrient limitation

The growth conditions for glucose and acetate must also be considered.

First, consider a batch cultivation in which all nutrients (including glucose and ammonium) are initially in great excess. As the culture proceeds, nutrients are gradually depleted, and the two microbes can reach a quasi-steady state together, as described in the previous section.

Alternatively, in a continuous culture such as a chemostat, all nutrients flow in and out at a dilution rate *D* [20]. *In this case, the effective loss rates become* 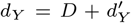 and 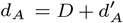, where 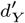 and 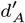 are the intrinsic death rates of yeast and acetobacter, respectively. The community growth rate is then *µ*_*C*_ = 0, with 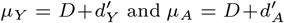. Both species must grow faster than the dilution rate to persist, and they will not grow at equal rates if their intrinsic death rates differ. A steady state is reached when all nutrient concentrations become constant.

Both species consume ammonium (as evident from their macrochemical equations in Figure 1), but this does not necessarily imply direct competition for ammonium. For example, if glucose is the limiting nutrient for the yeast and ethanol is limiting for the acetobacter, then their growth rates are determined by the concentrations of these respective energy sources. This occurs when the concentrations of glucose and ethanol are close to their Monod constants [21], while the ammonium concentration is much higher than its Monod constant and thus in excess. In such cases, the community is limited by the energy sources rather than by nitrogen availability.

### The consideration of maintenance requirements

Pirt’s interpretation of maintenance energy requirements for yeast and acetobacter can also be incorporated into the equations used to determine the relative abundances of the species [9].

We start from the steady-state requirement that the ethanol production flux equals the ethanol consumption flux:

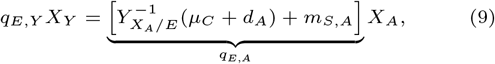

where *m*_*S,A*_ is the maintenance requirement of acetobacter, and 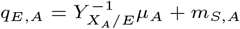 according to Pirt’s interpretation.

The maintenance requirement of yeast enters via the biomass yield of yeast on glucose, 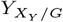

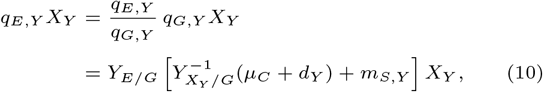

where *m*_*S,Y*_ is the maintenance requirement of yeast and 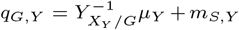 is Pirt’s equation for glucose uptake.

Combining these maintenance-extended expressions yields the following relation for the relative abundance of acetobacter:

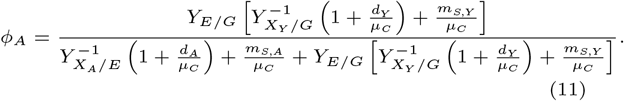

This expression reduces to equation 8 when the maintenance requirements are zero. Maintenance requirements influence relative abundances in a manner similar to death rates: increasing the maintenance requirement of a species decreases its relative abundance. Thus, incorporating maintenance does not qualitatively alter the insights obtained so far.

### The net conversion of the yeast–acetobacter community

At steady state, the net growth rates of yeast (*µ*_*Y*_ *− d*_*Y*_ ) and acetobacter (*µ*_*A*_ *− d*_*A*_) both equal the community growth rate *µ*_*C*_. The rates of the macrochemical reactions for yeast and acetobacter are therefore *µ*_*Y*_ *X*_*Y*_ and *µ*_*A*_*X*_*A*_, respectively.

For each reactant in the two macrochemical equations, the net production or consumption rate by the community equals the reaction rate multiplied by the stoichiometric coefficient of that reactant. For ethanol, these net rates cancel (produced by yeast, consumed by acetobacter). For other reactants, such as glucose, acetate, biomass, and ammonium, there is a net community-level production or consumption at steady state. The stoichiometry of the entire community at steady state is described by the macrochemical equation of the community, which we call the ‘community conversion’. This can be obtained directly from the macrochemical equations of the two species.

The derivation proceeds as follows. The two macrochemical equations occur at rates *µ*_*Y*_ *X*_*Y*_ and *µ*_*A*_*X*_*A*_. For example, the net consumption rate of ammonium by the community is:

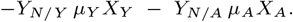

This rate must equal the rate of the community conversion (*µ*_*C*_ *X*_*T*_ ) multiplied by the stoichiometric coefficient of ammonium in that equation (*Y*_*N/C*_ ):

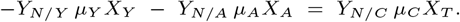

Thus, the ammonium coefficient in the community macrochemical equation is:

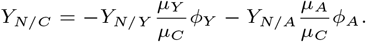

The same logic applies to all reactants in the net conversion equation. Cross-fed metabolites cancel out, and biomass terms require incorporating death rates.

In general, the macrochemical equation for the entire community (*MEQ*_*C*_ ) can be expressed in terms of the species-level macrochemical equations (*MEQ*_*Y*_ and *MEQ*_*A*_) as:

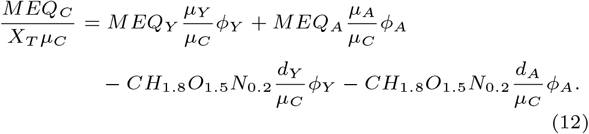

When the death rates are negligible, this reduces to the following macrochemical reaction for the community:

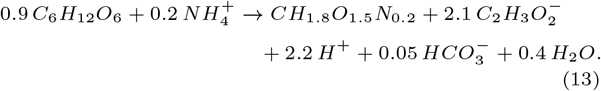

This equation constitutes the net conversion of the community—its “ecological service.”

A more general and compact derivation of the community net conversion, applicable to any community, is provided in the Appendix using a matrix formulation.

### Stoichiometric explanation of relative species abundances in terms of ATP metabolism

In the yeast–acetobacter community considered above, the biomass ratio is proportional to the product of two yield coefficients:

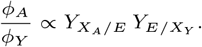

An increase in either coefficient changes the abundance ratio. For example, the yeast could increase 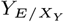 by producing more ethanol per unit biomass. This yield depends on the ATP requirement for biomass synthesis via fermentation: the ATP needed to assemble all macromolecules from central carbon metabolism precursors, minus the ATP produced during precursor biosynthesis. A more detailed treatment of this is given in the Appendix.

To generalise these ideas, consider a theoretical three-species community that sequentially converts a high-energy substrate *S*_1_ into a low-energy product *S*_4_:

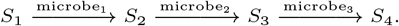

Here *S*_1_, *S*_2_, and *S*_3_ serve as both carbon and energy sources for the three species. For example, *S*_1_ could be glucose, *S*_2_ ethanol, *S*_3_ acetate, and *S*_4_ methane or carbon dioxide. Each substrate is partly converted into biomass and partly into the next catabolic product in the chain. Let *X*_1_, *X*_2_, and *X*_3_ denote the biomasses of the three species.

At steady state, the production and consumption rates of *S*_2_ and *S*_3_ must balance. This yields:

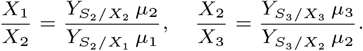

Here 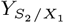 and 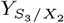 denote the amount of catabolic product formed per unit biomass, while 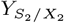 and 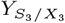 denote the amount of catabolic substrate required to form one unit of biomass.

All four yield coefficients can be linked to a common metabolic principle. A carbon–energy source (e.g., glucose) is used for:

1. Energy generation via ATP synthesis, and
2. Biosynthesis of precursors (e.g., pyruvate, glucose-6-phosphate) that are assembled into monomers for macromolecules (nucleic acids, amino acids, lipids).

Precursor biosynthesis can either generate ATP (e.g., pyruvate from glycolysis yields 2 ATP/glucose) or consume ATP (e.g., nucleotide synthesis from glucose-6-phosphate requires 1 ATP/glucose). The total ATP demand for biomass formation equals the ATP needed to convert precursors into macromolecules minus the ATP generated during precursor biosynthesis. The residual ATP demand must be met by catabolism, producing the next metabolite in the chain (*S*_2_ for microbe 1, *S*_3_ for microbe 2). The yields 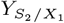 and 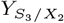 are therefore determined by this ATP balance, which in turn controls the steady-state abundances of *X*_2_ and *X*_3_.

In summary, the yield coefficients in macrochemical equations reflect the intracellular balance between catabolic ATP yields and anabolic ATP demands. Fermentation-product yields can be interpreted at coarse-grained level from this perspective – known as the “ATP method” for deriving macrochemical equations [25]. This framework directly links intracellular metabolism to relative species abundances in steady-state communities.

### Stoichiometric imbalance and spontaneous niche creation in simple cross-feeding communities

So far, we have only considered unidirectional cross-feeding. We now turn to bidirectional cross-feeding, where two species coexist and each produces a metabolite consumed by the other. Stoichiometric analysis predicts that, in general, one of these cross-fed metabolites will accumulate, creating a niche for a third species. We refer to this phenomenon as ‘stoichiometric imbalance’. Although it is easiest to illustrate with two species, the same reasoning applies to many-species communities.

Consider two species, *A* and *B*: species *A* produces metabolite *a* and consumes metabolite *b*; species *B* produces *b* and consumes *a*. In addition to cross-feeding, we allow for outflow of both *a* and *b* from the system. At steady state, the metabolite balances are:

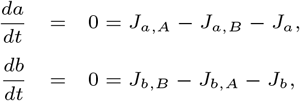

where *J*_*a,A*_ is the production rate of *a* by species *A, J*_*a,B*_ is the consumption rate of *a* by species *B*, and *J*_*a*_ is the outflow rate of *a* (similarly for *b*). Using the approach from previous sections, these fluxes can be expressed in terms of yields, growth rates, and biomass:

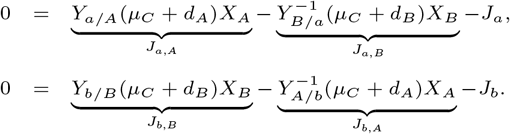

Defining *X*_*T*_ = *X*_*A*_ + *X*_*B*_, *ϕ*_*A*_ = *X*_*A*_*/X*_*T*_, and *ϕ*_*B*_ = *X*_*B*_*/X*_*T*_, the *a*-balance gives:

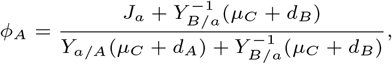

while the *b*-balance gives:

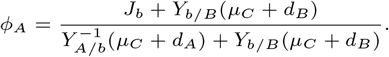

In steady state, these two expressions for *ϕ*_*A*_ must be equal. Two cases arise:

- **Stoichiometrically balanced:** *J*_*a*_ = *J*_*b*_, which can include *J*_*a*_ = *J*_*b*_ = 0. In this special case, the elemental composition of *a* and *b* and the metabolic demands of *A* and *B* align perfectly, so neither metabolite accumulates. This is expected to be rare.
- **Stoichiometrically imbalanced:** *J*_*a*_ ≠ *J*_*b*_, which we expect to be the general case. Here, one metabolite accumulates relative to the other.

In the imbalanced case, natural selection is expected to drive the community toward maximal growth. Under this condition, one overflow rate (*J*_*a*_ or *J*_*b*_) will approach zero. If *J*_*a*_ = 0, the system is *A*-limited, *J*_*b*_ *>* 0, and a niche exists for a third microbe that can grow on *b*. Conversely, if *J*_*b*_ = 0, the system is *B*-limited and a niche exists for a microbe that can grow on *a*.

In summary, stoichiometric reasoning alone predicts that bidirectional cross-feeding typically leads to the accumulation of one cross-fed metabolite, thereby creating a new ecological niche.

### The carrying capacity and the net metabolic conversion of a large community

In figure 2, a community consisting of five microbial species is shown. Together they convert glucose (e.g. deriving from plant litter) into carbon dioxide, methane and biomass. The species feed on each other’s waste products, via cross feeding of acetate, hydrogen, butyrate and carbon dioxide. A representative macrochemical reaction of each of the microbial species is shown and were obtained from Smeaton & van Capellen [26].

**Fig. 2.**
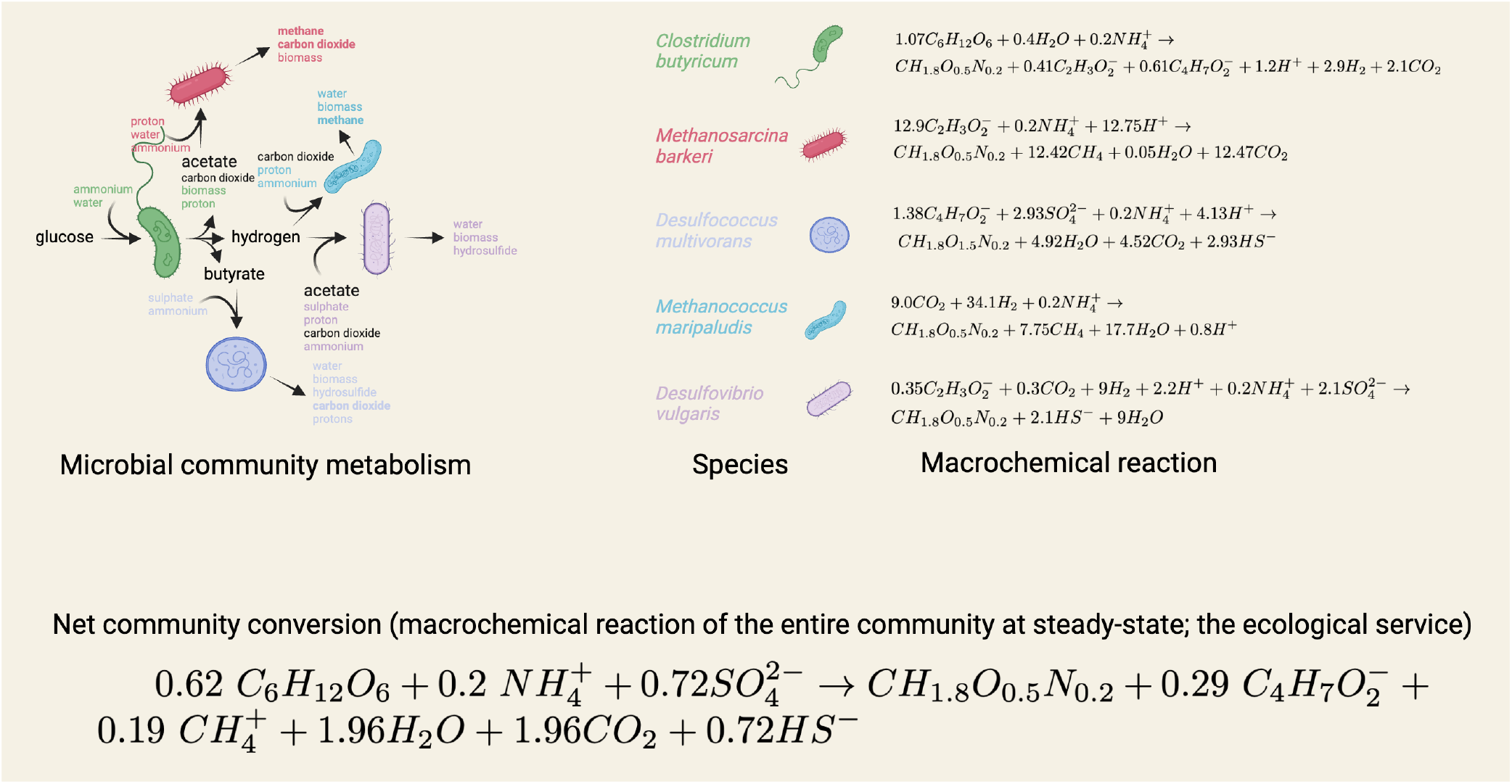
The metabolism and ecological service of a community of five microorganisms in terms of their macrochemical equations. The macrochemical equations of five microorganisms are shown that together convert glucose into methane, biomass and carbon dioxide, via the metabolic intermediates: acetate, hydrogen, butyrate and acetate. This net metabolic conversion of the microbial community, it’s ecological service, is shown as a macrochemical equation of the community. The ecological service can be computed from the relative abundances of the species and the macrochemical equations of the community using the method outlined in the main text. Chemical compounds: *C*_6_*H*_12_*O*_6_, (glucose), 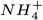(ammonium), *CH*_1.8_*O*_0.5_*N*_0.2_ (biomass), *CH*_4_ (methane), *H*_2_*O* (water), *CO*_2_ (carbon dioxide), 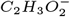(acetate), 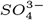(sulphate), *HS*^*−*^ (hydrosulfide).

In the Appendix we provide a complete stoichiometric analysis of the microbial community shown in figure 2. We show how the species fraction can be expressed in terms of the stoichiometric coefficients of the macrochemical equations of the species, the net fluxes of nutrient in and out of the system, the community growth rate, and the death rates of all the species. This summarises all the methods in a matrix formalism that is applicable to more complex cases then those considered so far. This analysis also involves a numerical example, which leads to the following community conversion at steady state: glucose, ammonium and sulphate are converted into biomass, butyrate, method, carbon dioxide, and hydrosulfide,

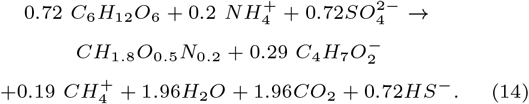

In this case, four of the five species *C. butyricum, D. vulgaris, D. multivorans*, and *M. barkberi* are active at, respectively, the following biomass fractions:

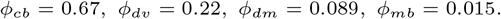

Glucose functions here as the energy source and 0.72 mole of it is needed to make 1 gram biomass. The carrying capacity of the environment equals therefore 1/1.9 mole biomass per mole glucose. This solution was obtained using community flux balance analysis [27] – since more fluxes occur in the community than stead-state flux balances. We note that since the community makes butyrate (and methane) a niche still exists for a butyrate (and methane) consumer to fill.

## Discussion

It seems that we are still far from a quantitative understanding of microbial communities such that we can predict and explain why particular species are carrying out specific metabolisms. A first step in that direction is understanding the relations between the metabolic stoichiometry of microbial growth (macrochemical equations), the relative abundance of species, and the net metabolic conversion of a steady-state community. Here we presented this understanding and provided various extensions such as the maintenance requirement, living and dead biomass, stoichiometric imbalance, and various community topologies (incl. those analysed in the Appendix).

Beyond providing a conceptual understanding, the balance equations derived here can be applied directly to experimental data. For example, relative abundances can be inferred from measured fluxes, especially when combined with biomass measurements and stable isotope probing (SIP) proteomics [28, 29]. SIP proteomics can help link specific metabolic activities to individual species, and the framework can be integrated with community flux balance analysis in cases where the system of equations is underdetermined due to limited data. Many models in microbial ecology, such as the generalised Lotka Volterra or the resource-consumer models, do not consider stoichiometry. These phenomenological models are particularly useful for communities of poorly characterised species or when the focus is on abundance dynamics rather than precise metabolic activities – for instance, the dynamics of their abundances in terms of generalised species-species interactions [30]. Cases also exist where the interest is primarily in terms of the consequences of the metabolic capacities and interactions between the species. For example, how they concertedly maintain a particular metabolic ecological service such as, for instance, the anaerobic conversion of plant litter into greenhouse gases [6]. Then stoichiometric models are more relevant than more phenomenological models.

Stoichiometric models can describe certain phenomena better than non-stoichiometric models. A straightforward example occurs for the exchange of two nutrients *a* and *b* between two species *A* and *B. A* makes *a* and feeds on *b* made by *B*, which feeds on *a*. (For instance, *a* is acetate and *b* is an amino acid.) In such a cross-feeding exchange it will generally be so that the system settles to a steady state with equal specific growth rates of the two species and that one of the exchanged nutrients remains constant by a balanced production and consumption, while the other accumulates (stoichiometric imbalance; see results section). This accumulation can not be predicted by a non-stoichiometric model of the microbial community. Whether *a* or *b* accumulates depends on the precise stoichiometry of *A*’s and *B*’s macrochemical equation. If *a* accumulates, then other species can invade the community then when *b* accumulates. Stoichiometric models are needed to distinguish between these scenarios.

Finally, although we accounted for the presence of living and dead biomass, we did not model the recycling of dead biomass components. This extension was omitted because the stoichiometry of biomass recycling is currently too poorly understood, but it could be incorporated once better quantitative information becomes available [23].

## Acknowledgements

We thank Francesco Moro, Matti Gralka and Jack Pronk for discussions about some of the material presented in this paper.

## Derivation of the macrochemical reaction of a steady-state metabolic network

Consider a metabolic network with stoichiometric matrix **N** at steady state such that its flux vector **J** obeys

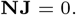

The macrochemical equation (MEQ) that corresponds to this state of the metabolic network can be obtained from

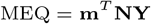

with: **m** is the vector with metabolite names (ordered according to the row ordering of the stoichiometric matrix) and **Y** as the vector that results after dividing all entries of **J** by the value of the growth rate (often one of the entries of **J**). The MEQ can be written in its macrochemical reaction form by adding all the negative terms left of a chemical reaction arrow and those that are positive to its right-hand side.

## Derivation of equations appearing in the main text

Derivation of equation 1

Given *ϕ*_*i*_(*t*) = *X*_*i*_*/X*_*T*_, we obtain

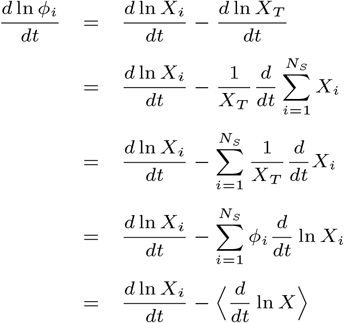

With 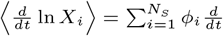 ln *X*_*i*_, i.e. the average *d* ln *X*_*i*_*/dt*.

Derivation of equation 16

The relative abundances *L*_*i*_*/X*_*T*_, *D*_*i*_*/X*_*T*_ and *X*_*i*_*/X*_*T*_ = *L*_*i*_*/X*_*T*_ + *D*_*i*_*/X*_*T*_ are all fixed at any steady state of the community. This implies the following relations:

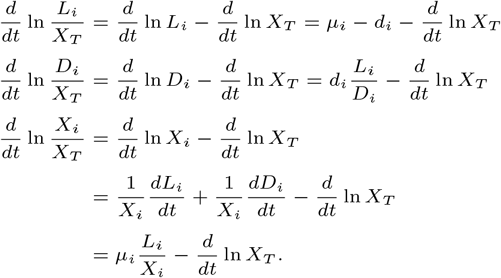

Hence, the following relations hold for the community growth rate equals

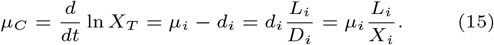

From these relations, we can deduce that

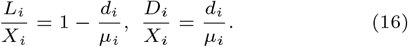

These fractions sum to 1, since *X*_*i*_ is defined as the sum of *L*_*i*_ + *D*_*i*_.

We already showed that the community growth rate *µ*_*C*_ equals *d* ln *X*_*T*_ */dt* and the mean value of *d* ln *X*_*i*_*/dt*. An expression of this mean can be derived,

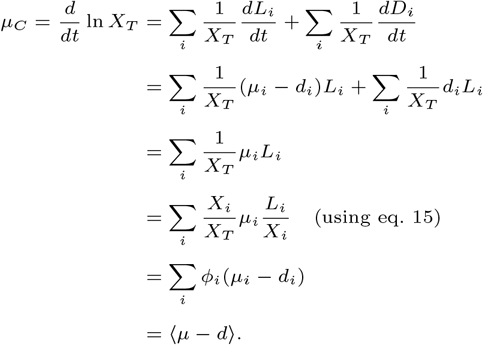

Thus, this last result and equation 15 indicate that the growth rate of each species equals, at any steady state of the community,

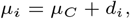

which implies that the concentrations of the extracellular nutrients and products – of which are toxins for other species – which impact the growth rate of species are eventually such that this relation holds. E.g. in the simplest case the growth rate of a species depends only on the concentration of a single limiting nutrient concentration according to a Monod equation. Since the metabolic activity of a species is proportional to its growth rate and its energy-maintenance rate, the growth rate sets the net uptake and production rates of nutrients and products of one particular species. If a microbe cannot achieve the community growth rate it goes extinct.

Finally, using *µ*_*C*_ = *µ*_*i*_ *− d*_*i*_ and equation 16, we conclude that

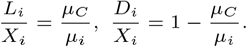

## Derivation of the catabolic product and biomass yield expression in terms of ATP yields

We limit ourselves to a carbon source as energy source, from which a catabolic product is made as well as precursor molecules, consumed during the synthesis of amino acids, nucleic acids, and lipids.

Consider the following simplified representation of a 2-species community

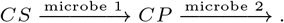

*CP* is the catabolic product of microbe 1, made from its carbon source *CS. CP* is also the carbon source for microbe 2.

At steady state of the community, the net production and consumption rate of the catabolic product balances

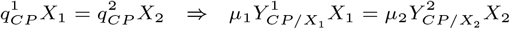

with 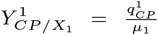 as the yield of catabolic product on biomass of microbe 1 (both made from *CS*) and 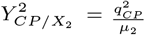, which is best represented as the inverse yield of biomass of microbe 2 on the catabolic product of microbe 1, i.e.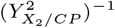. These yields are net stoichiometric properties of the entire metabolic networks of these two microbes, are stoichiometric coefficients of the macrochemical reactions of the two species, and are each calculable with flux balance analysis from a genome-scale stoichiometric model of each of the species, from experiments or from phenomenological models.

The last equation shows that

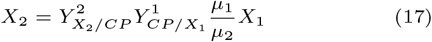

and that therefore the biomass abundance of the second microbe depends on the yield of catabolic product on biomass of microbe 1 and the biomass yield of microbe 2 on this same compound. Next, it will be shown that both yield can be written in the ATP metabolism of those two microbial species.

Next, we will express the yield of catabolic product on biomass 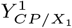 and the yield of biomass on a catabolic substrate 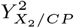 in terms of the ratio of the amount of ATP produced during catabolic-product formation over the amount produced during precursor metabolism of both species.

The total specific carbon source flux uptake rate *q*_*CS*_, in terms of *CS* for microbe 1 and in terms of *CP* for microbe 2, equals the sum of the carbon flux towards catabolic product (*CP* ) and precursor (*Pre*) metabolism,

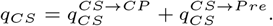

Precursor metabolism involves the synthesis of metabolites from the carbon source that act as precursor for the biosynthesis of lipids, nucleic acids and amino acids. For instance, alanine is made from the precursor metabobolite pyruvate on glucose growth and yields 1 ATP per pyruvate, while the glucose-6-phosphate, a precursor for nucleic acid biosynthesis via the pentose phosphate pathway, requires 1 ATP per glucose-6-phosphate. When a precursor is made then no catabolic product is made, e.g. no ethanol, acetate, or formate, results from pyruvate when it is used as precursor. Catabolic-product formation is always accompanied by a net gain in ATP. The amount of catabolic product made depends on the ATP requirement that remains after subtraction of the ATP requirement per unit biomass by the amount made during precursor biosynthesis (this amount can be negative).

The ratio of carbon flow towards precursors versus catabolic product is defined by the following ratio of specific flux values,

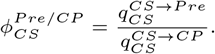

Both microbes synthesize their ATP as part of the conversion of carbon source to catabolic product (*CS → CP* ) and during precursor formation from carbon source (*CS → Pre*),

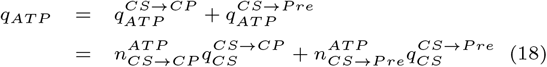

The specific production rate of catabolic product is defined by,

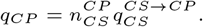

The ‘n’ stoichiometric coefficients are properties of the metabolic pathways that two microbes use to make catabolic products and precursor metabolites from catabolic substrates. Next, we need to determine 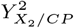, which is a yield of biomass of microbe 2 on its carbon source,

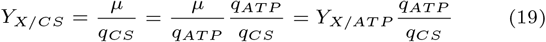

We assume that we know the yield of biomass on ATP (i.e. *Y*_*X/AT P*_, which is also quite conserved across microbes), and quantifies how much ATP is needed to make 1 unit of biomass. Finally, we determine the yield of ATP per unit carbon source 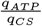 in terms of the ratio of ATP made during precursor biosynthesis over that made during catabolic product formation,

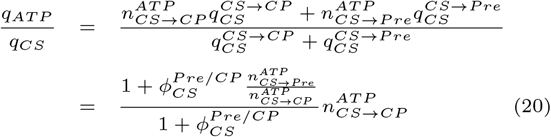

We still need to determine the yield of catabolic product on biomass, i.e. for microbe 1: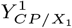. The yield of a catabolic production of biomass can also be expressed in terms of whole-cell ATP stoichiometries,

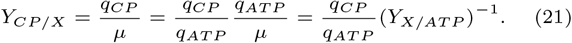

Next, we express 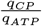 in terms of the stoichiometric coefficients of intracellular metabolism,

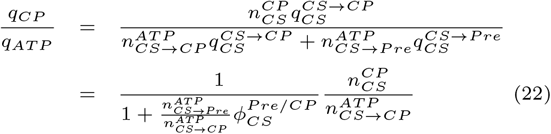

With those four relations (eq. 19, 20, 21, and 22), we can express the ratio of biomass abundances in terms of the *ATP* - stoichiometries of the metabolism of the two species, by their substitution in equation 17,

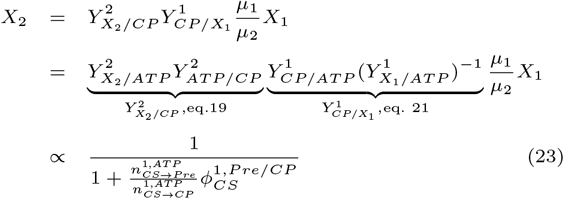

What becomes clear is that when microbe 1 meets its ATP demand for macromolecule biosynthesis from precursor molecules predominantly via precursor biosynthesis, hence 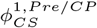 is high, then it does not make much CP per unit biomass and, hence, the relative biomass abundance of species 2 is low. Thus, biomass abundance of the second species depends on the ATP stoichiometry of microbe 1 that makes the catabolic substrate of the microbe 2.

## Illustration of the calculation of relative species abundances and community conversion

### Calculation of the relative species abundances of a steady state community

The relative species abundances are determined from the steady-state condition of the metabolites that are exchanged between the species, which are also allowed to flow in or out of the community. We will take anaerobic digestion of glucose by a microbial community (shown in figure 2) as an example.

This community has four exchanged (variable) metabolites (i.e. acetate (ac), carbon dioxide (co), butyrate (bu) and hydrogen (h2); we consider glucose fixed) and five species (*Clostridium butyricum* (cb), *Methanosarcina barkeri* (mb), *Desulfococcus multivorans* (dm), *Methanococcus maripaludis* (mm), and *Desulfovibrio vulgaris* (dv)).

Since we have four variable metabolites, we have four flux balance equations that hold at steady state,

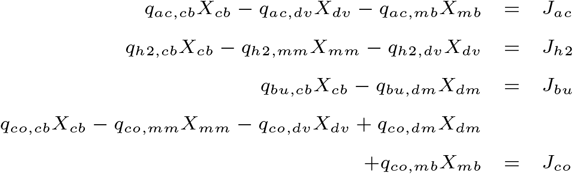

The term on the left are the production and consumption rates of the species while the fluxes on the left (i.e. the *J*’s) are the exchange fluxes with the environment. If glucose would be considered flowing into the environment, instead of fixed as it is now, then one more balance for glucose would have added.

Next, we introduce yields and relative species abundances. The yields are introduced by multiplying and dividing each term in the previous equations by the associated growth rate of the species, e.g.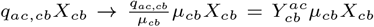. The relative species abundances are introduced by division of all terms by the total biomass abundance *X*_*T*_, i.e.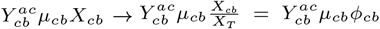. The resulting set of equation can be couched in a matrix equation to which we also add the conservation relation for the species fractions.

This equation can be simplified, by including the condition that the growth rate of each species equals the community growth rate (*µ*_*C*_ ) plus the death rate of the species, e.g. *µ*_*cb*_ = *µ*_*C*_ + *d*_*cb*_. We assume that we either know the growth rate of each of species or the community growth rate and the death rate of each species. From the macrochemical equations we obtain the yield coefficients. This allows for the determination of the species fractions by solving the previous set of linear equations

### Calculation of the community conversion (the ecological service) of a steady state community

To determine the community conversion – the macrochemical equation – of the steady-state community, we start from the steady-state flux vector **f** of the community,

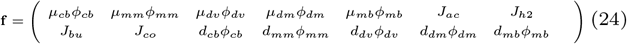

We also require the names of all the nutrient concentrations and species abundances occuring in the system. These we collect in a vector ordered in the same order as the rows of the stoichiometric matrix. Thus vector we refer to as **n**,

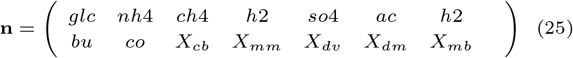

We also need to matrix **S** containing the stoichiometric coefficients of all the processes that occur in the community.

The following multiplication then gives the macrochemical equation *MEQ* of the microbial community or the net conversion of the community.

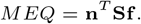

### Numerical example anaerobic digestion

The numerical example of the anaerobic digestion community and leading to the macrochemical equation, shown in figure 2, corresponds to the following case. The relative abundances are:

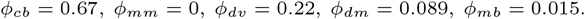

The fluxes are: *J*_*glc,i*_ = 1, *J*_*nh*4,*i*_ = 0.28, *J*_*bu,o*_ = 0.40, *J*_*ch*4,*o*_ = 0.26, and *J*_*CO*2,*o*_ = 2.73, 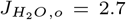, *J*_*HS,o*_ = 1 and *J*_*SO*4,*i*_ = 1 (with ‘i’ denoting inflow and ‘o’ outtflow out of the community). This solution was obtained with linear programming (maximisation of the growth rate of the community).

## Additional theoretical examples

### The carrying capacity and the net metabolic conversion for a 2-species community

The description of a two species community with the exchange of a single metabolite, similar to the yeast and acetobacter community, can be done in general terms using two macrochemical reactions and two death processes as shown in Figure 3. For simplicity, we only focus on the reactants that matter for the example (the nutrients *A, B* and *C* (e.g. glucose, ethanol and acetate) and the biomasses *X*_1_ and *X*_2_). Those could for instance be referring to the carbon (and energy) sources and products and do not consider a maintenance requirement.

**Fig. 3.**
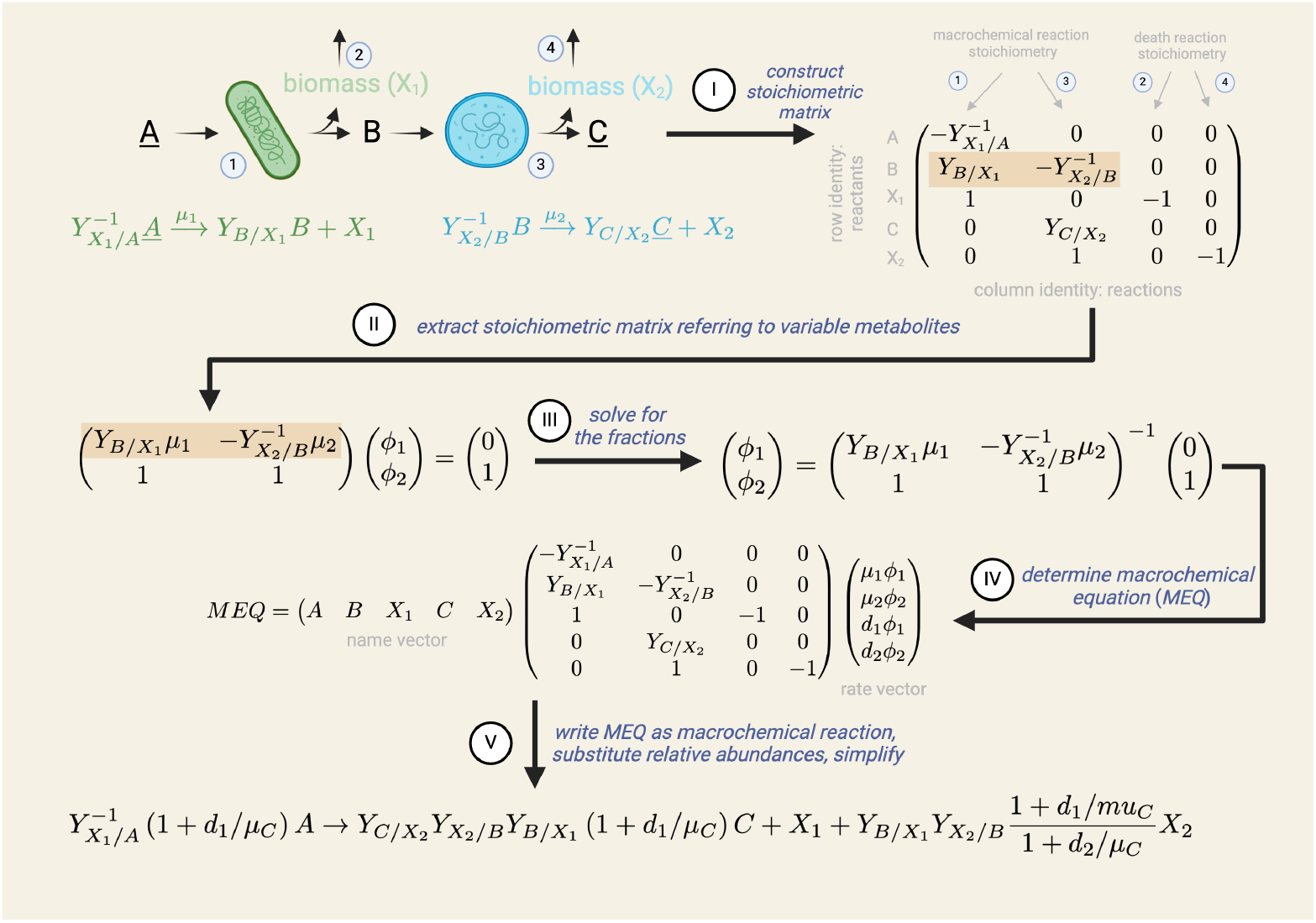
Illustration of the derivation of the macrochemical reaction and carrying capacity of a 2-species community. The underline indicates that we consider the concentration of the nutrients A and C to be fixed, to allow for the occurrence of a steady state. We assume that *B* does not overflow into the environment.

**Fig. 4.**
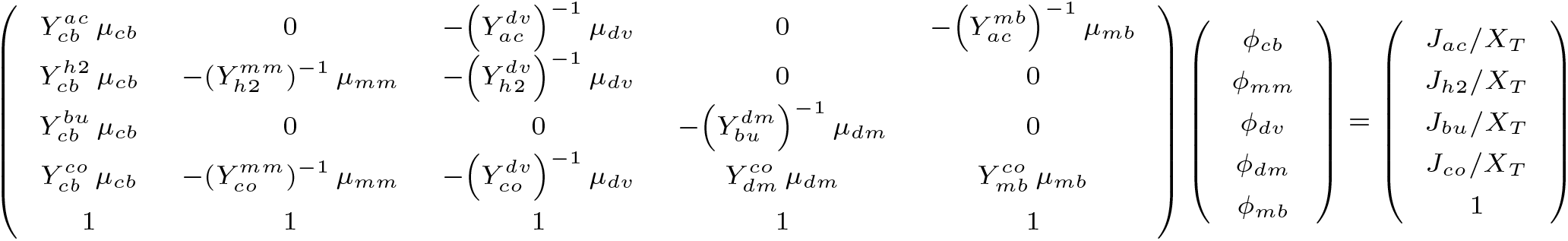
Community mixing relation for species fractions *ϕ* with exchange fluxes *J*_*•*_ (per *X*_*T*_ ).

**Fig. 5.**
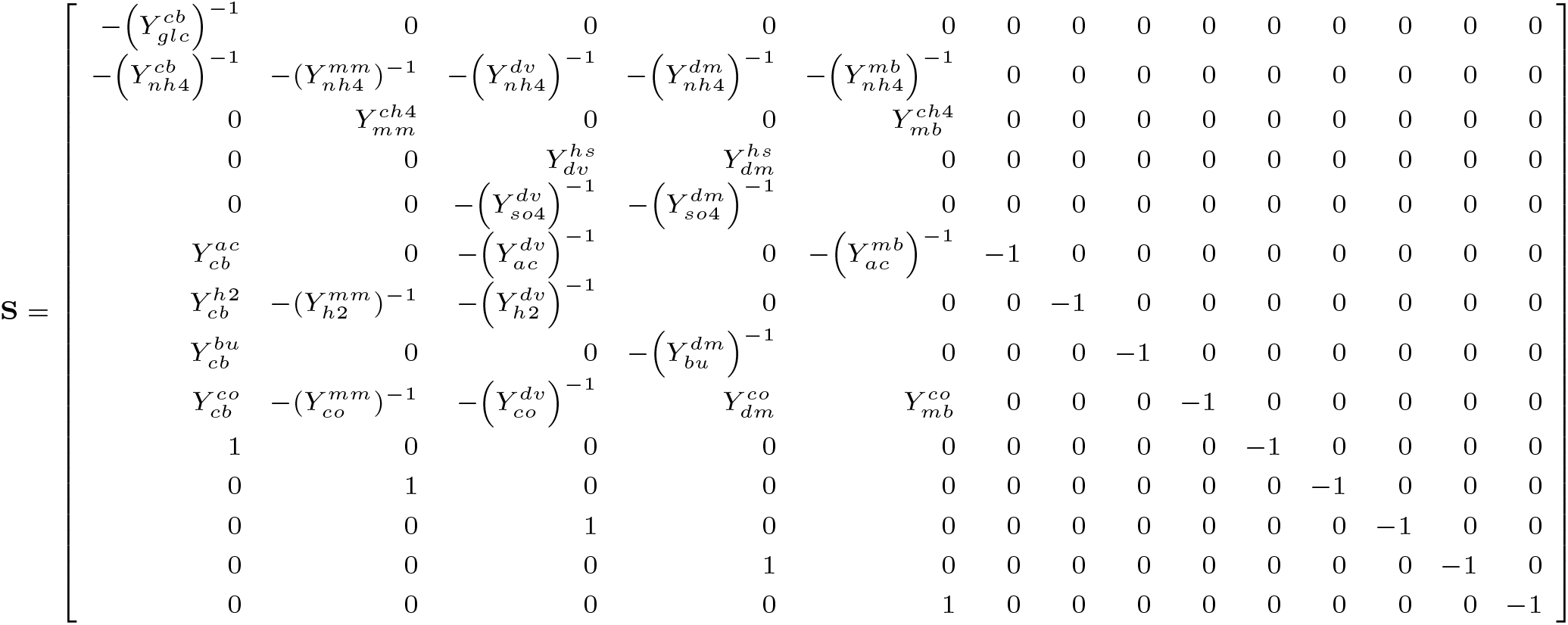
Stoichiometric matrix **S** for the community processes.

In figure 3, the stepwise procedure for the determination of the community conversion of a community is shown, starting from the macrochemical reactions of its species.

The first step is to construct the stoichiometric matrix of the metabolism of the community. This matrix has four columns corresponding to the two growth and two death reactions and five rows, i.e. the three nutrients and the two biomasses. The derivation of the community conversion requires first the derivation of the species fraction (step 2). This is done from the steady-state balances for the concentration of the variable nutrients in the network and the conservation relationship for the fraction. In this case we have only one balance equation, i.e. for *B*. After the determination of the fractions, the community conversion is obtained by multiplying the stoichiometric matrix determined in step 1 from the left by the name vector of all the species and from the right by the steady-state flux vector. This equation can then be written as a reaction by gathering all the terms with a negative sign left of the reaction arrow and those with a positive sign to its right. In this way we obtain to obtain the stoichiometry of the net conversion of the community per unit *X*_1_,

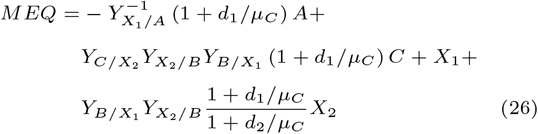

We consider this macrochemical reaction as the ecological service of the microbial community. This can, for instance, be the recycling of a chemical element or the conversion of plant litter into methane and carbon dioxide, which we work out in the Appendix.The community conversion (eq. 26) indicates that *A* is consumed to make *C, X*_1_ and *X*_2_. When the death rate of the first microbe would increase more *A* is consumed to produce more *C* and *X*_2_ per unit *X*_1_; so the ratio of *X*_2_*/X*_1_ increases. The carrying capacity of the community equals the yield coefficients in front of the biomasses (divided by the yield coefficient of *A* in case one would like to express the capacity per unit of the limiting nutrient).

When the community as a whole is not growing in size, i.e. *µ*_*C*_ = 0, all microbes have a growth rate that equals their death rate. The macrochemical equation of the community then simplifies to

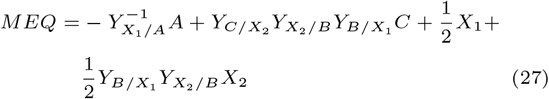

The community conversion will remain constant as long as none of the two species changes its metabolism in such a way that its macrochemical equation and net growth rate changes.

### Branched microbial communities and branch flux ratios

Many microbial communities are branched and two or more species consume the same nutrient, which is the byproduct of the growth process of (an)other species. An example is shown in Figure 1B. A branched network can be treated in a similar way as above, but since we end up with fewer exchanged metabolites between species than the number of species we need to add more information. We will explicitly consider a small example to make this more clear. (The general case is treated in the Appendix in terms of some linear algebra properties of the stoichiometric matrix.)

We consider the following branched network composed out of 4 species and 2 exchanged nutrients,

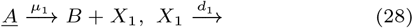

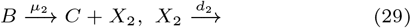

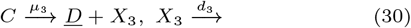

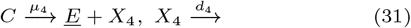

The following concentration balances exist for the two metabolic intermediates

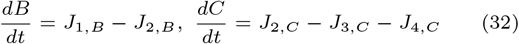

Leading to the the following equations at steady state (using the same method outlined above),

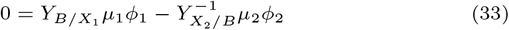

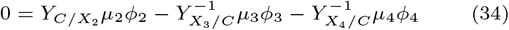

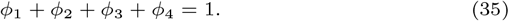

We aim to determine the four relative abundances. We are however one equation short (we have 4 *ϕ*’s but only 3 equations that relate them) and require an additional relation between relative abundances to determine the relative abundances of the species. If we know (by measurement) the community’s synthesis rate of *D* and *E*, then we know their ratio and obtain a new relation between relative abundances, i.e.

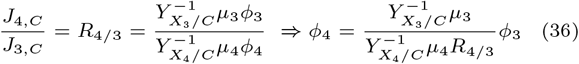

This extra relation for the determination of all *ϕ*’s form a set of linear equations. The associated equations are shown in the Appendix.

The net conversion of the community can again be calculated using the method explained in the previous section. It depends on the measured flux ratio and the ratio’s of death rates and growth rates.

### Cyclic microbial communities, chemical element conservation and a cycle law

Since many microbial communities conversions are associated with the recycling of chemical elements, we also consider a community that has a cyclic exchange of nutrients between species, driven by an external thermodynamic driving force. We consider a 4-species cycle where the first organism feeds on *A*, which is recycled by the fourth microbe,

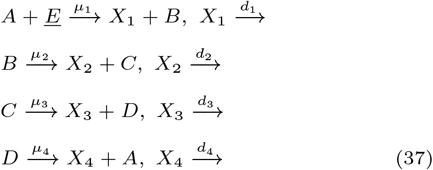

As before, we do not show the other nutrients occurring in the macrochemical reaction which we assume fixed. In this case, we have four exchanged nutrients and species, corresponding to the following concentration balances,

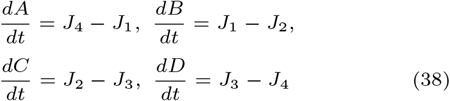

Since the sum of these four equations equals 0, we have to conclude that the following sum of concentrations is conserved such that at all times the following relation holds: *T* = *A* + *B* + *C* + *D*.

Due to this linear dependency between the concentration balances, we only have three independent steady-state linear relations for the fractional abundances and their summation to 1,

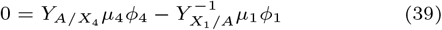

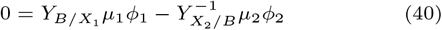

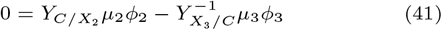

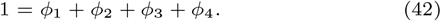

This set of equations is solvable and gives rise to the following abundance ratios

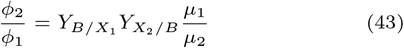

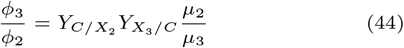

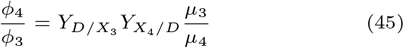

We can also deduce that since 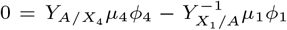 the following relation holds as well

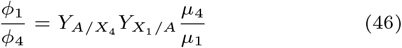

Since this equation needs to be in agreement with the previous abundance ratios, we deduce from the previous derived ratios that

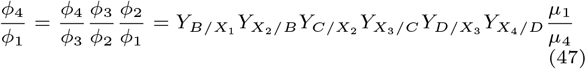

and that therefore the following relations holds for the product of the yields along the cycle,

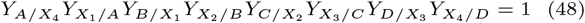

To understand how it arose, consider the implication of this equation that (upon cancellation of all growth rates and multiplication of the denominator and numerator by the product of the biomasses)

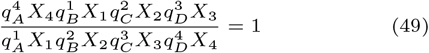

which is correct since this equals *J*_1_*J*_2_*J*_3_*J*_4_*/*(*J*_1_*J*_2_*J*_3_*J*_4_) = 1. This equation is reminiscent of the detailed balance equations of that exist for cycles in chemical reaction networks which state that in thermodynamic equilibrium the net free energy change is zero.

## Author contributions statement

Must include all authors, identified by initials, for example: S.R. and D.A. conceived the experiment(s), S.R. conducted the experiment(s), S.R. and D.A. analysed the results. S.R. and D.A. wrote and reviewed the manuscript.

